# Microbial community changes correlate with impaired host fitness of *Aurelia aurita* after environmental challenge

**DOI:** 10.1101/2023.03.01.530242

**Authors:** Nicole Pinnow, Cynthia M. Chibani, Simon Güllert, Nancy Weiland-Bräuer

## Abstract

Climate change globally endangers certain marine species, but at the same time, such changes may promote species that can tolerate and adapt to varying environmental conditions. Such acclimatization can be accompanied or possibly even be enabled by a host’s microbiome; however, few studies have so far directly addressed this process. Here we show that acute, individual rises in seawater temperature and salinity to sub-lethal levels diminished host fitness of the benthic *Aurelia aurita* polyp, demonstrated by up to 34 % reduced survival rate, shrinking of the animals, and almost halted asexual reproduction. Changes in the fitness of the polyps to environmental stressors coincided with microbiome changes, mainly within the phyla Proteobacteria and Bacteroidota. The absence of bacteria amplified these effects, pointing to the crucial importance of a balanced microbiota to cope with a changing environment. In a future ocean scenario, mimicked by a combined but milder rise of temperature and salinity, the fitness of polyps was severely less impaired, together with condition-specific changes in the microbiome composition. Our results show that the effects on host fitness correlate with the strength of environmental stress, while salt-conveyed thermotolerance might be involved. Further, a specific, balanced microbiome of *A. aurita* polyps is essential for the host’s acclimatization. Microbiomes may provide a means for acclimatization, and microbiome flexibility can be a fundamental strategy for marine animals to adapt to future ocean scenarios and maintain biodiversity and ecosystem functioning.

## 1. Introduction

It is widely recognized that marine ecosystems are under threat (Consortium 2021). Ocean acidification, the global increase in sea surface temperature, and change in salinity along with overfishing, eutrophication, sedimentation, and pollution endanger marine species globally (Buck-Wiese, Voolstra & Brüwer 2016; Ducklow *et al*. 2022; Leal Filho *et al*. 2022). At the same time, marine organisms adjust and cope with changing environments (Fuhrman 2009; Reusch 2014; Foo & Byrne 2016; Abirami *et al*. 2021). Eco-physiological studies typically focus on solitary macroorganisms or interactions among them (Sutherland *et al*. 2013; Martini *et al*. 2021), such as competition (e.g., acidification influencing turf algae-kelp interactions) and predation (e.g., warming leading to kelp grazing by range-expanding herbivorous fishes) (Vergés *et al*. 2016; Provost *et al*. 2017; Qiu *et al*. 2019).

In contrast, less is known about the impact of ocean climate change on interactions between macro- and microorganisms (Harvell *et al*. 2002; Hutchins & Fu 2017; Hutchins *et al*. 2019). All multicellular organisms live in an intimate and interdependent association with their microbiome that includes bacteria, archaea, viruses, fungi, and protists (Bosch & McFall-Ngai 2011; Faure, Simon & Heulin 2018; Jaspers *et al*. 2019; Hentschel 2021). Consequently, animals and plants represent functional biological entities comprising a host and its microbiome, so-called metaorganisms (Bosch & McFall-Ngai 2011). Members of a host-associated microbiota have various functions within a metaorganism and display fundamental roles in host health by contributing, for instance, to host development (Rook *et al*. 2017), organ morphogenesis (Sommer & Bäckhed 2013), metabolism (Ochsenkühn *et al*. 2017; Shaffer *et al*. 2017), aging (Gould *et al*. 2018), behavior (Ezenwa *et al*. 2012), and reproduction (Chilton *et al*. 2015; Jacob *et al*. 2015; Weiland-Bräuer *et al*. 2020a). Microbes may further be essential for macroorganisms living in extreme environmental conditions (Bang *et al*. 2018) and for acclimating and adapting to environmental changes (Torda *et al*. 2017; Ziegler *et al*. 2017; Pita *et al*. 2018; Voolstra & Ziegler 2020; Baldassarre *et al*. 2022). Although the host can also respond to environmental perturbations through phenotypic plasticity (Yampolsky, Schaer & Ebert 2014; Foo & Byrne 2016; Pazzaglia *et al*. 2021), microbial-mediated acclimatization has received particular attention. Microorganisms have shorter generation times and respond more rapidly and versatilely than their hosts (Peixoto *et al*. 2021).

In Nature, metaorganisms face a diversity of biotic and abiotic stressors that may require an associated microbial community that responds adequately (Carrier & Reitzel 2017). Accordingly, a metaorganism can maximize fitness through compositional changes in the associated microbiota (abundance and/or diversity) (Rosenberg & Zilber-Rosenberg 2014; Bordenstein & Theis 2015; Bang *et al*. 2018; Voolstra & Ziegler 2020; Peixoto *et al*. 2021). Such dynamic restructuring of a host’s community through environmental change is known as microbiome flexibility. For instance, microbiome flexibility has been proposed to play a role in the rapid acclimatization of *Fungia granulosa* after long-term exposure to high-salinity levels (Röthig *et al*. 2015), acclimatization of *Acropora hyacinthus* to increased thermal stress (Ziegler *et al*. 2017), and the ability of the coral and sponge holobiont to cope with environmental change (Ribes *et al*. 2016; Ziegler *et al*. 2019; Voolstra & Ziegler 2020). If novel traits and functions in the microbiome can significantly affect the metaorganism phenotype to enable acclimatization, such traits may be vertically transmitted to the next generation and ultimately lead to metaorganism adaptation (Webster & Reusch 2017; Bang *et al*. 2018). However, only a few studies have directly addressed how a microbiome enables acclimatization to short-term changes in a local environment or enables adaptation of the host (e.g., (Moran 2007; Soen 2014; O’Brien *et al*. 2019)).

To provide insights into these processes, our research is focused on the microbiome of the moon jellyfish *Aurelia aurita* (Linnaeus) and its involvement in the eco-physiological responses of that host. The scyphozoan *A. aurita* is a cosmopolitan species documented worldwide in various coastal and shelf sea environments (Dong 2019) and is also one of the most frequent blooming jellyfish species (Pikesley *et al*. 2014; Dong 2019). The biphasic life cycle of *Aurelia* alternates between free-living pelagic medusae that sexually reproduce and generate planula larvae that form sessile benthic polyps (**Fig. 1A**). Polyps can undergo asexual reproduction through daughter polyp generation by budding and ephyrae (precursor medusae) constriction during strobilation (Lucas 2001). Environmental factors such as temperature, salinity, or food supply influence both the asexual reproduction of the polyps and medusa ecology, such as somatic growth and sexual maturation (Pitt & Kingsford 2000; Xing *et al*. 2020; Schäfer *et al*. 2021).

**Fig. 1:**
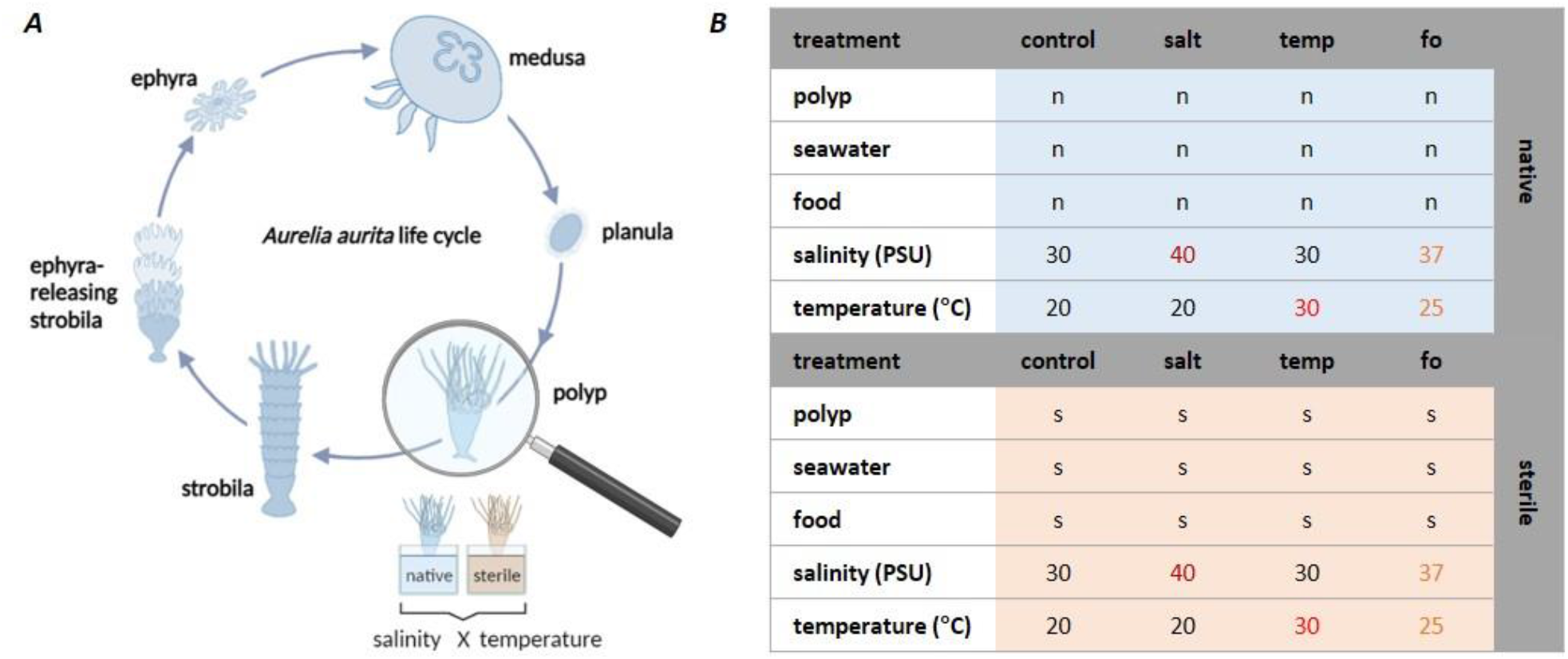
Study design of the host-fitness experiment. **(*A*)** The life cycle of *Aurelia aurita* alternates between pelagic medusae and benthic polyps. The host-fitness experiments were conducted with polyps exposed to increased temperature and salinity. **(*B*)** Each treatment comprised 96 native (n) or sterile (s) polyps (the latter were kept under sterile conditions throughout the experiment). Control conditions included a salinity of 30 PSU and an ambient temperature of 20 °C. Salinity was increased to 40 PSU (salt) or 37 PSU (fo); temperature was raised to 30 °C (temp) or 25 PSU (fo).

*A. aurita* is a highly flexible species that can adapt to a wide range of environmental conditions and can survive and reproduce between 4 – 28 °C and 15 – 38 PSU salinity (Bamstedt 1990; Widmer 2005; Pascual *et al*. 2015; Widmer, Fox & Brierley 2016). Temperature plays a crucial role in the reproduction of polyps (e.g., (Willcox, Moltschaniwskyj & Crawford 2007; Purcell *et al*. 2012; Loveridge, Lucas & Pitt 2021; Schäfer *et al*. 2021). At higher temperatures, polyps tend to reproduce daughter polyps by budding, while below a certain threshold, strobilation is triggered to reproduce planktonic ephyrae (Schäfer *et al*. 2021). Salinity is expected to determine the settlement of planulae and subsequent development of polyps (Conley & Uye 2015; Dong 2019) and may also affect the distribution of polyps in coastal waters (e.g., (Watanabe 2001; Willcox, Moltschaniwskyj & Crawford 2007; Sokołowski *et al*. 2016), and the mortality of polyps (Janas & Witek 1993; Sokołowski *et al*. 2016). Understanding the effect of abiotic factors on the survival and reproduction of *A. aurita* is essential for accurate predictions on the species’ future under climate change and its potential to bloom: jellyfish blooms significantly impact an ecosystem’s composition and structure by changing element cycling and interfering with food webs (Dong 2019; Goldstein & Steiner 2019).

The composition and structure of the microbial communities associated with *A. aurita* are well characterized and was shown to be crucial for *A. aurita’s* fitness (survival, feeding, and growth) and particularly for the generation of offspring (Weiland-Bräuer *et al*. 2020a). Here, we tested how *Aurelia*’s microbiome is changing in composition due to acute temperature and salinity rises, thus affecting host fitness. Ultimately, the microbiome of this metaorganism might mediate the acclimatization of *A. aurita* to climate change, and as a first step in this process, short-term changes were investigated here.

## 2. Materials and Methods

### 2.1 *Aurelia aurita* polyp husbandry and generation of sterile polyps

Husbandry and generation of sterile polyps are described in detail in previous studies by Weiland-Bräuer *et al*. (Weiland-Bräuer *et al*. 2015; Weiland-Bräuer *et al*. 2020a; Weiland-Bräuer *et al*. 2020b). Briefly, polyps of sub-population North Atlantic (Roscoff, France) were kept in the laboratory at 20 °C in 30 PSU artificial seawater (ASW containing 3 % w/v Tropical Sea Salts, Tropic Marin) and fed twice a week with freshly hatched *Artemia salina* (HOBBY, Grafschaft-Gelsdorf). These conditions simulate the mean sea surface temperature in summer (20 °C) and salinity (30 PSU) of the North Atlantic Ocean, where these polyps originated (Gac *et al*. 2021). Sterile polyps and *A. salina* were generated with Provasoli’s antibiotic mixture (360,000 U/L penicillin G, 1.5 mg/L chloramphenicol, 1.8 mg/L neomycin, and 9,000 U/L polymyxin B; Carl Roth, Karlsruhe, Germany). The absence of bacteria was confirmed by the lack of amplification of the bacterial 16S rRNA gene with a standard PCR using primer set 27F and 1492R (Lane 1991). Sterile animals received sterile food.

### 2.2. Challenge of *Aurelia aurita* polyps with environmental stressors

Host fitness experiments were conducted according to Weiland-Bräuer *et al*., 2020 (Weiland-Bräuer *et al*. 2020a), with a similar setup for native and sterile conditions (**Fig. 1**). Applied conditions were kept constant throughout the experiments. Single native or sterile polyps were transferred from husbandry tanks to 48-well plates. Each well containing a single polyp was filled with 1 mL native or sterile ASW (filtered through 0.22 µm filters). All treatments were conducted with 96 replicates each (**Fig. 1B**). Native and sterile polyps were exposed to control conditions (20 °C and 30 PSU) and raised temperatures (30 °C, 30 PSU) or increased salinity (40 PSU, 20 °C). A future ocean scenario was simulated by combining 37 PSU ASW with 25 °C (**Fig. 1B**). The latter values were based on a predicted increase of 5 °C and 2 PSU in the year 2500 under the assumption of a 0.1 °C increase per decade and a total salinity rise of 5 % (Cheng *et al*. 2020; Jo *et al*. 2022). Even at present, heat waves can cause relatively abrupt temperature and salinity anomalies within this range (Feudale & Shukla 2011a; Feudale & Shukla 2011b; Miyama, Minobe & Goto 2021). The experimental conditions were maintained for four weeks.

### 2.3 Monitoring host-fitness traits

Six different fitness traits: survival, growth, feeding, budding, strobilation, and ephyrae release were monitored. All animals were recorded over time using a stereomicroscope (Novex Binokulares RZB-PL Zoom-Mikroskop 65.500, Novex, Arnhem, the Netherlands) equipped with an HDMI/HD camera. The survival of polyps was assessed every 48 h for the first 14 d based on their phenotypical appearance and the presence of tentacles, and accumulative death was calculated at day 14. Growth was documented every 48 h during the same period by measuring the length and width of the polyps. Mean start sizes (length multiplied by width at t_0_) and after 14 d (t_14_) were compared per treatment, and growth rates (in %) were calculated. Budding was monitored by counting the number of daughter polyps, and the weekly budding rate was calculated for the first 14 days. The feeding rate of the polyps was monitored for five days during week 3 of the experiments. For this, single polyps were offered 20 *Artemia salina*, and after 1 h, the remaining prey was counted; a mean feeding rate (% of Artemia clearance) was then calculated per treatment. Strobilation and ephyrae release were monitored in parallel. For this set of experiments, strobilation was induced by adding 5 μM 5-methoxy-2-methyl indole to the water at days 1, 2 and 3 (involving daily washing and inductor exchange as described in (Weiland-Bräuer *et al*. 2020a)). On day 4, polyps were washed with water to remove the inducer. Strobilation was monitored daily, and strobila phenotypes and the number of segments were detected beginning on day 5 when native control polyps began segmentation. Ephyrae release was monitored each day after their first appearance, and the number of released ephyrae was detected beginning on day 12. Ephyrae release was monitored for the next 4 weeks.

### 2.4 Data analysis of host-fitness parameters

For each treatment, fitness trait parameters were analyzed, resulting in the following fitness variables: (i) survival, calculated from counts of alive and dead polyps, (ii) % growth rate, (iii) % feeding rate, (iv) % budding rate, (v) counts of segments and the number of ephyrae (strobilation). Fitness variables were assessed using univariate permutational analysis of variance (Anderson 2001). All fitness variables were tested in the 10 most informative pairwise comparisons between the twelve treatments. PERMANOVAs were performed using R. The vegan package was used for the computations, and the permutation test for adonis was performed under the reduced model with 9,999 permutations (Dixon 2003; Oksanen *et al*. 2013).

### 2.5 16S rRNA amplicon-based microbiota analysis

16S rRNA amplicon sequencing was performed to analyze the microbial community composition of native polyps. Six native polyps were randomly removed from the 96 replicates after 14 d. DNA isolation and subsequent 16S rRNA amplicon sequencing were performed as previously described (Weiland-Bräuer *et al*. 2020a)). DNA was isolated using the WIZARD Genomic DNA Purification kit (Promega, Madison, WI, USA), and PCR amplicon libraries of the V1-V2 region of the 16S rRNA gene were constructed using uniquely barcoded primers with primers V1_A_Pyro_27F (5’
s-CGTATCGCCTCCCTCGCGCCATCAGTCAGAGTTTGATCCTGGCTCAG-3’) and V2_B_Pyro_27F (5’-CTATGCGCCTTGCCAGCCCGCTCAGTCAGAGTTTGATCCTGGCTCAG-3’) combined with 338R. Following amplification in 20 μL, the amplicons were sequenced on an Illumina MiSeq v3 platform (2 × 300 cycle kit) at the Max-Planck Institute for Evolutionary Biology in cooperation with Dr. S. Künzel. 16S rRNA data processing was conducted with mothur v1.39.5 (Schloss *et al*. 2009) according to the MiSeq SOP (OTUs were detected at a 97% similarity threshold) using SILVA SSU database 138 as described in (Weiland-Bräuer *et al*. 2020a). All downstream computations were performed in R v4.0.0. Sequence data were deposited under the NCBI BioProject PRJNA925707, and BioSample Accessions SAMN32807491-SAMN32807530.

## 3. Results

Host-fitness experiments were conducted with *A. aurita* polyps with a high number of replicates (N=96) to elucidate the effect of temperature and salinity rises on this host and decipher its microbiota’s role for any acclimatization potential. The host fitness traits of survival, growth, feeding and asexual reproduction were studied under various combinations of increased temperature and salinity (**Fig. 1**), as these environmental stress conditions are linked to climate change. Treatments of native polyps harboring a diverse but stable microbial community under the applied artificial lab conditions were compared with identical treatments of sterile polyps that had been generated with antibiotics.

### 3.1 Increased temperature and salinity affect host survival, growth, and feeding rates

The phenotype of native polyps exposed to control conditions, when their survival rate was 100 %, is shown in **Fig. 2A**. When the salinity was increased to 40 PSU or the temperature to 30 °C, polyp survival was significantly reduced (**Fig. 2B**). The effect of raised salinity on survival was not prominent, giving a 9 % reduction, but this was still significant (p < 0.001, the outcome of all PERMANOVA tests on survival is summarized in **Tab. S1**). Polyps exposed to high salinity developed a thickened polyp body and absorbance of tentacles (**Fig. 2A**, salt treatment). The elevated temperature resulted in a 34 % reduction of the survival rate (p-value < 0.001), and live polyps frequently developed an impaired phenotype with absorbed tentacles and a roundish body shape (**Fig. 2A**, temperature treatment). The combination of increased salinity and temperature in a future ocean scenario (fo treatment) produced slight, non-significant effects on polyp survival (99 %, p = 0.265; **Fig. 2B**). Note that the elevation of the single parameters was more extreme (30 vs. 25 °C and 40 vs. 30 PSU) than applied in the combination simulating a future ocean. The observed differences in survival might depend on the extent of the increase in temperature and salinity or on a salt-conveyed thermotolerance. Although their survival was unaffected, the polyps in conditions resembling a future ocean appeared abnormal, with thickened and round body shapes (**Fig. 2A**).

**Fig. 2:**
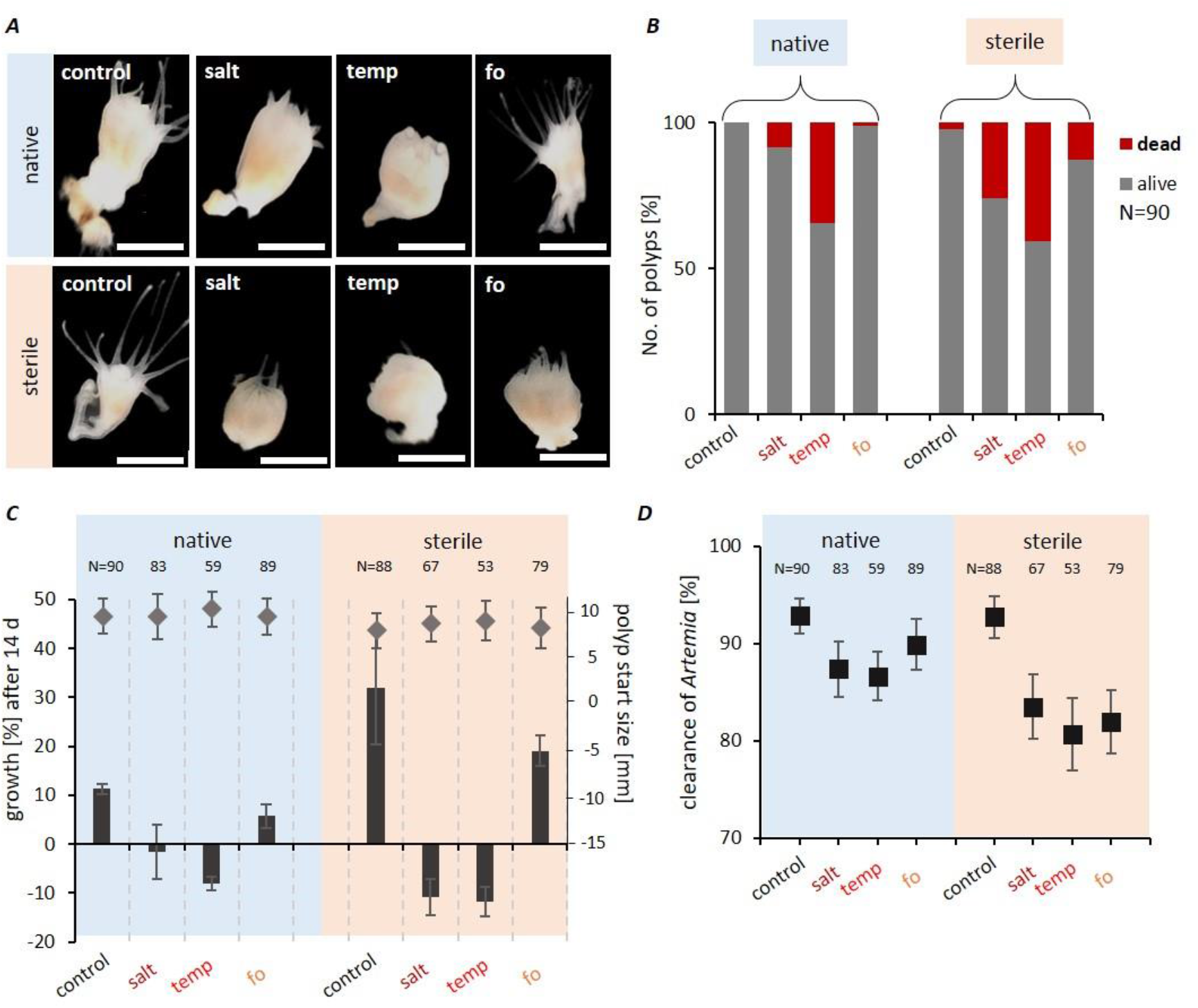
High temperature and salinity impair the fitness of *Aurelia aurita* polyps. **(*A*)** Photographs of typical polyps for each treatment after 14 d. Scale bars correspond to 1 mm. **(*B*)** Percentages of dead and alive polyps (90 biological replicates) based on polyps’ phenotypical appearance and presence of tentacles. Monitoring was conducted each 48 h, and accumulative numbers are shown after 14 d. **(*C*)** The growth of polyps was followed every 48 h for 14 d by measuring the polyp size (length times width) of live polyps. Mean start sizes (♦, legend at the right) and the corresponding growth rates (%, legend at the left) are plotted. **(*D*)** Feeding rate (% clearance of 20 *Artemia salina* in 1 h) plotted as mean of single polyps of five monitoring days.

Similar experiments were also performed with sterile polyps. High-temperature stress lowered their survival rate to 59 % (**Fig. 2B**). Thus, the absence of a microbiome decreased the survival of temperature-stressed animals by a further 8 % compared to native polyps (p = 0.023). In the future ocean scenario, the survival rate of sterile polyps was decreased to 87 %, which was lower than native animals kept under these conditions (p = 0.027 native vs. sterile in fo, **Tab. S1**). Overall, the survival trends observed with native polyps were exacerbated in sterile animals.

The treated polyps were analyzed for growth after 14 d (**Fig. 2C**). Under normal conditions, native polyps had a mean size (length multiplied by width) of 9.6 ± 2.08 mm^2^ at the beginning of the experiment, and this increased to 10.6 ± 2.4 mm^2^ after 14 days, corresponding to a growth rate of 11 % (**Fig. 2C**). Exposure to higher salinity halted growth after 14 days (**Fig. 2C**, p < 0.001; the outcome of PERMANOVA tests for growth is summarized in **Tab. S2**), while high temperatures resulted in downsized polyps (**Fig. 2C**, p-value < 0.001). Under future ocean conditions, the polyps could grow, albeit significantly less than under control conditions (growth rate 6 %, **Fig. 2C**, p < 0.001). Sterile animals were slightly smaller than native animals at the start, but when the growth of sterile polyps was permitted, their growth rates were generally higher compared to native animals (**Fig. 2C**). Growth was not permitted in the absence of microbes under high salinity or high temperature. The shrinking effect under salt stress was substantial, where sterile polyps decreased their size by 9.3 % compared to halted growth of native animals (**Fig. 2C**, p < 0.001). Furthermore, the sterile animals shrank more strongly at high temperatures than the native polyps (−11.8 % vs. -8.0 %, p = 0.012).

Feeding rates of live polyps were assessed during week three of the experiments for five consecutive days (**Fig. 2D**). Native polyps kept under control conditions had a mean *Artemia* clearance rate of 92.8 % ± 8.9 % (**Fig. 2D**). High salinity or high temperature caused a reduction to 87.4 % and 86.6 %, respectively (**Fig. 2D**, p = 0.03 and p = 0.036; see **Tab. S3** for PERMANOVA tests on feeding). The feeding rate of sterile polyps was significantly reduced (p < 0.02) under all stress conditions compared to the control treatment of those animals (**Fig. 2D**). There was no statistical difference in feeding rates between native and sterile animals kept under the same conditions (**Tab. S3**). Thus, the increased growth of sterile animals compared to native polyps under the same condition was not due to increased feeding.

### 3.2 Increased salinity and temperature affect the asexual reproduction of the host

To determine the effect of the environmental stressors on the asexual reproduction of *A. aurita*, the generation of daughter polyps was monitored every 48 h over 14 days (**Fig. 3A**). Under control conditions, native polyps showed an average budding rate of 0.15 daughter polyps per week (**Fig. 3B**). They generated up to 2 daughter polyps per week (9 % produced one daughter polyp, and 3 % resulted in two daughter polyps). Budding of native polyps was significantly negatively affected in all stress treatments (**Fig. 3A**, B; p < 0.001, see **Tab. S4** for PERMANOVA tests on budding). Under all stress conditions, fewer polyps were produced that lacked tentacles (example photographs are shown for the fo condition in **Fig. 3A**). In the absence of bacteria, budding was seriously impaired. Under control conditions, the budding of sterile polyps was decreased by 45 % (**Fig. 3B**, p < 0.001, **Tab. S4**), in line with previous observations (Weiland-Bräuer *et al*. 2020a). Only 8 % of the sterile animals produced one daughter polyp, and none produced two. The applied environmental stress conditions lowered the reproduction rates even further (**Fig. 3D**, p < 0.001 for all sterile conditions compared to sterile control treatment, **Tab. S4**); no daughters were produced at all by sterile polyps under high temperature (**Fig. 3A, B**).

**Fig. 3:**
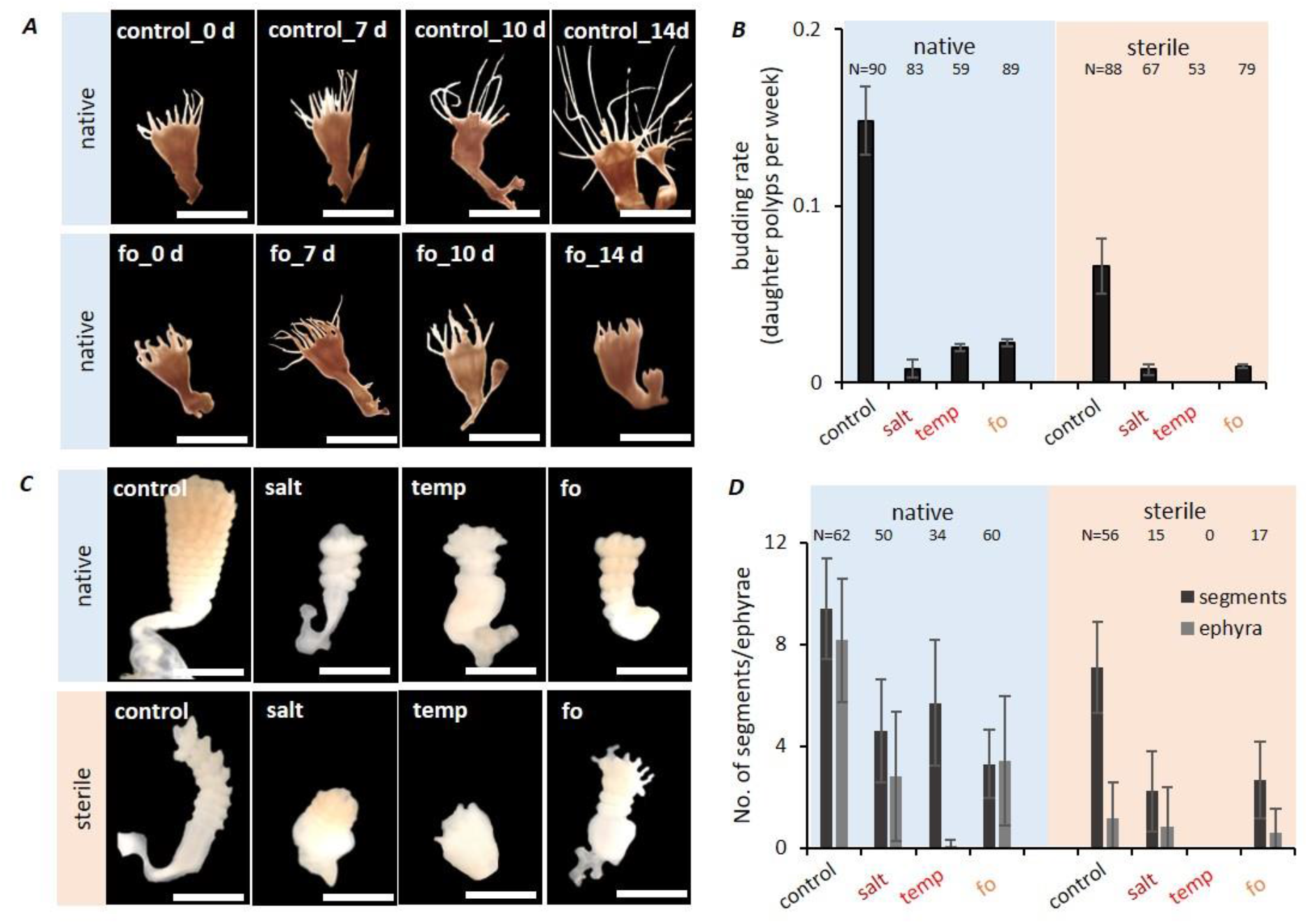
Asexual reproduction of *Aurelia aurita* polyps diminished under ambient stress conditions. **(*A*)** Daughter polyp generation of native polyps under control and future ocean conditions followed for 14 d. Scale bars correspond to 2 mm. **(*B*)** Budding was followed every 48 h for 14 d by monitoring the generation of daughter polyps and calculated as daughter polyp generation per week. **(*C*)** Strobilation of polyps was induced using 5 µM 5-methoxy-2-methyl indole. Photographs show the segmentation of polyps for native and sterile polyps under control conditions six days post-induction. Scale bars correspond to 2 mm. **(*D*)** Formation of strobilae was monitored daily and quantified by the number of segments formed per strobila. The numbers of released ephyrae were determined for 4 weeks.

Following chemical induction, native polyps kept under control conditions began strobilation by initiating the segmentation of the prolonged body (early strobila) on day 5. Here, segmentation was completed in the late strobila stage on day 9 (**Fig. 3C**), and by then, 62 of the 96 replicate native polyps (65 %) had formed a strobila (indicated by the number of strobilae above **Fig. 3D**). Raising the salinity and temperature significantly reduced the formation of strobilae, which were now only visible in 50 and 34 polyps, respectively (52 % and 35 %, a significant reduction compared to control p < 0.001, **Tab. S5**). The number of segments formed per polyp under the various treatments on day 9 is shown in **Fig. 3D**. All stress conditions of native polyps resulted in fewer numbers of segments (p < 0.001, **Tab. S5**), and subsequently, fewer ephyrae were released by native animals under stress conditions (**Fig. 3D**, p < 0.001, **Tab. S6**). The formation of strobilae was not only impaired but also delayed, and abnormal phenotypes were formed (**Fig. 3C**), including malformed structures, incomplete or completely missing constriction, impaired tentacle absorption, and colorless, minimized, and partly thickened bodies (**Fig. 3C**). Compared to control conditions, in which native polyps constricted a mean of 9.4 segments to release eight ephyrae (**Fig. 3D**), environmentally stressed polyps formed only three to five segments and released between zero and 3.4 ephyrae (**Fig. 3D**). The most potent effect was observed at high temperature (6 segments, no ephyrae, **Fig. 3D**; p < 0.001). The elevated temperature and salinity levels, alone or in combination as in the future ocean scenario, produced even more substantial negative effects on asexual reproduction without the microbiota. Crucially malformed strobilae were monitored, showing only slight constrictions, which went hand in hand with massively reduced ephyrae release (0 to 0.8; p < 0.001). The offspring’s generation was halted entirely at raised temperatures without a microbiota (**Fig. 3C, D**).

### 3.3 Environmental stress conditions cause changes in microbial community patterns that correlate with reduced host fitness

Six native polyps were randomly taken from the 96 replicates for each treatment after 14 d, and 16S rDNA V1-V2 amplicon sequencing was performed to characterize the polyp microbiota. A subset of 2,300 sequences per sample was generated to eliminate bias due to unequal sampling. In total, 946 OTUs were identified, and of these, 461 OTUs were shared by all biological replicates of all treatments. Phylogenetic analysis of the samples revealed a complex microbiota structure that changed due to exposure of the polyps to environmental stressors (**Fig. 4**). The reproducibility between the six replicates per treatment was high, and thus, the means of the replicates were reported. At the phylum level, the microbiome of polyps kept under control conditions was composed of 74 % Proteobacteria and 18 % Bacteroidota, with 4 % reads derived from Firmicutes and 3 % unclassified (uncl.) bacteria, whereas bacterial lineages with < 1 % relative abundance (collectively reported as “others”) accounted for 1 % (**Fig. 4A**). Within the phylum Proteobacteria, *Vibrio* and *Alteromonas* accounted for the largest proportion; Bacteroidota were mainly represented by *Ulvibacter* and uncl. Bacteroidota. Although Proteobacteria, Bacteroidota, Firmicutes, and uncl. Bacteria remained the most abundant phyla also under stress conditions, shifts were observed (**Fig. 4A**). The higher salinity resulted in a stark reduction in Proteobacteria (now only comprising 38 %) in favor of Bacteroidota (now 53 %), while Firmicutes and uncl. Bacteria remained almost constant. Although not quite as intense, similar shifts were observed at high temperatures. Here, 52 % Proteobacteria and 34 % Bacteroidota were assigned, while Firmicutes were detected with 2 % and uncl. Bacteria with 6 % (**Fig. 4A**). In comparison, the future ocean treatment showed weaker changes on phylum level compared to control conditions, giving 70 % Proteobacteria, 24 % Bacteroidota, 1 % Firmicutes, 2 % uncl. Bacteria, and 3 % others (**Fig. 4A**). Major shifts at the genus level are summarized in **Fig. 4B**. For all stress treatments, an increase of at least 1 % was observed for uncl. Sinobacteraceae, uncl. Saprospiraceae and uncl. Flavobacteriaceae, at the expense of *Alteromonas, Pseudoalteromonas, Pseudomonas*, uncl. Proteobacteria and *Exiguobacterium* (**Fig. 4B**).

**Fig. 4:**
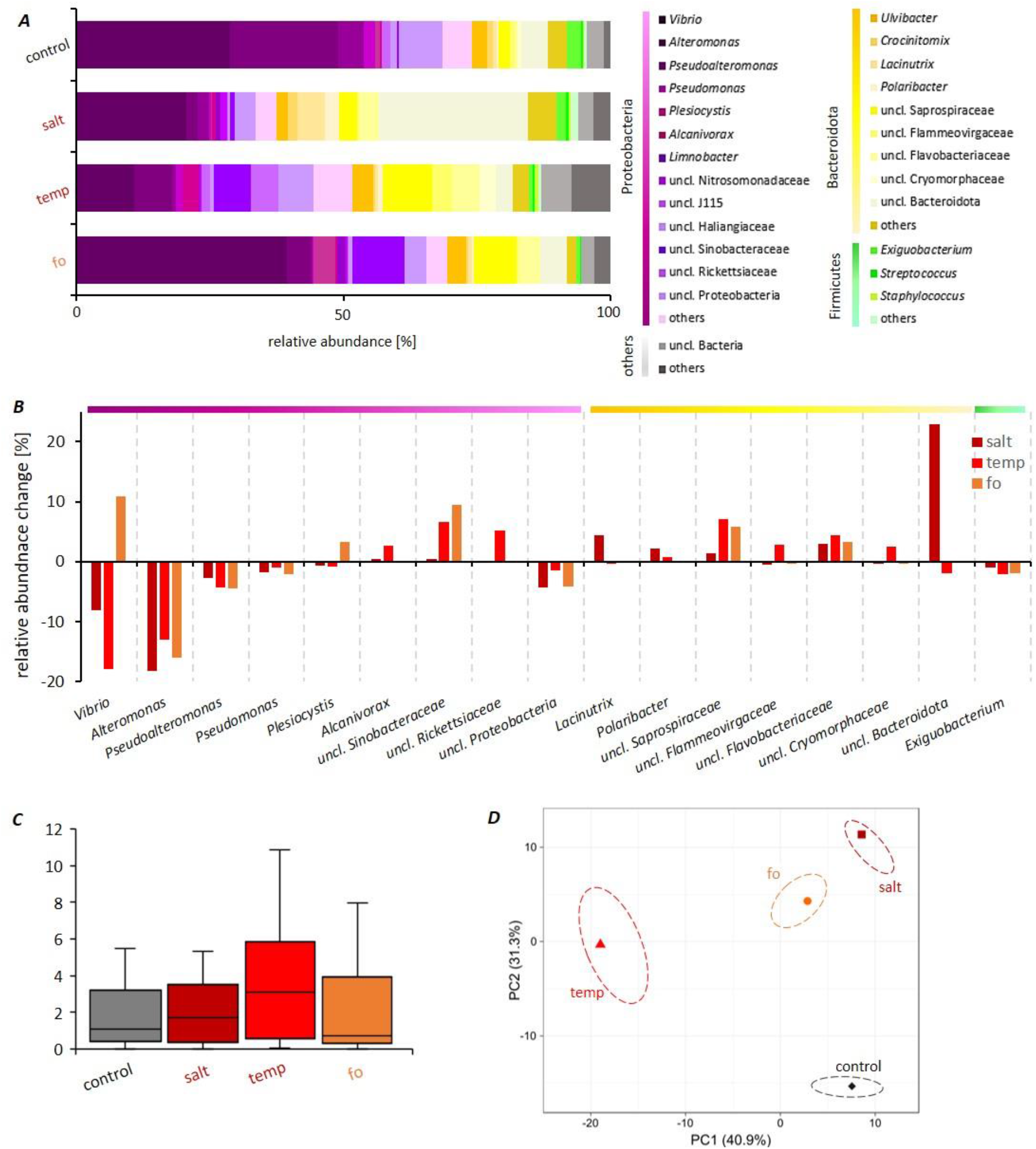
Microbial community composition of *Aurelia aurita* polyps under environmental challenge. Microbial communities were analyzed by sequencing the V1-V2 region of the bacterial 16S rRNA gene. OTU abundances were summarized at the genus level and normalized by the total number of reads per sample. **(*A*)** Bar plots visualizing the dominant genera (reaching at least 1 % of relative abundance) after 14 days of maintenance under control and at the indicated conditions. All data are based on the means of 6 biological replicates. **(*B*)** Differences in relative abundances at day 14 compared to control conditions for all dominant genera. **(*C*)** Boxplot of alpha-diversity Shannon indices. **(*D*)** Principal component analysis (PCA) for genus-level microbiota of the polyps (mean of 6 replicates). Different colors and shapes denote treatments and ellipses representing 95 % confidence interval for the centroids of each data cluster.

*Vibrio* showed a decline in salt and temperature treatments, whereas an increase was recorded for the future ocean scenario (**Fig. 4B**). *Alcanivorax*, uncl. Rickettsiaceae, uncl. Flammeovirgaceae, and uncl. Cryomorphaceae increased in relative abundance exclusively after temperature increase (**Fig. 4B**), while *Lacinutrix* and, in particular, uncl. Bacteroidota proliferated under salt stress (**Fig. 4B**). Despite these differences, the alpha-diversity (species richness and evenness as calculated by the Shannon index) of the polyps’ microbiota did not significantly differ between treatments. However, the range between replicates was notably smaller in the salt treatment compared to the two other treatments (**Fig. 4C**). Beta-diversity was assessed by Principle component analysis (PCA) at the genus level to summarize the marked differences in community composition of the individual polyps (**Fig. 4D**). The first two axes of the generated PCA plot explain 72.2 % of the variation of the analyzed communities, that were separated into four clusters corresponding to the treatments. The microbiota resulting from high temperature produced the highest variance in bacterial composition compared to control conditions. High salinity and the mild but combined heat- and saline stress of the future ocean had less impact on the compositional variance, suggesting that the variance between the bacterial compositions depends on the strength and the applied environmental stress, whereby the effects differ for each applied condition (**Fig. 4D**).

Hierarchical cluster analysis verified the different community structure profiles obtained from all conditions and resolved the observed microbiota dynamics on the OTU level. The first analysis considered all shared 461 OTUs and identified their changes in relative abundances compared to the control treatment. These were grouped taxonomically in the heatmap, visualizing their fold-change (**Fig. 5A**). Shifts can be seen across the whole microbial community; however, 26 % of the OTUs (122 of 461) remained constant (cutoff 0.2 % relative abundance change, p < 0.0001) compared to the control conditions. The combination of salinity and temperature stress in the future ocean scenario not only resulted in specific changes but also reflected the effects of separate elevated temperatures or salinity. We next zoomed in on the 31 most abundant OTUs (relative abundance > 1 % in the control), as their shifts explained the major differences in the community composition after the applied stresses (**Fig. 5B**). By far, the strongest effects were seen for OTU 001 (*Alteromonas*, strongly decreasing in all three treatments) and OTU 0010, an unclassified Bacteroidota that strongly increased with high salt. Eight OTUs decreased mildly in abundance to all conditions (OTUs 0019, 0156, 0148, 0014, 0060, 0050, 0101, 0017 in cluster VIII) and OTUs 0029 and 0001 decreased more strongly, while 21 OTUs increased to most of the stressors (**Fig. 5B**). However, only OTU 0122 (an uncl. Rickettsiaceae member of the Proteobacteria) remained constant under salt and fo conditions compared to the control. Of the increased OTUs, twelve proliferated under all stress conditions (OTUs 0003, 0007, 0093, 0013, 0015, 0028, 0024, 0030, 0047, 0023, 0109, and 0033). An increase due to future ocean conditions but a decrease in relative abundance through individual salt and temperature treatment was detected for OTUs 0002 and 0006. OTUs 0039 and 0005 were raised due to salt stress but declined under temperature and fo conditions. Two OTUs (0085 and 0057) increased with temperature while decreasing under salt and fo treatments. Only OTU 0004 increased under temperature and fo conditions, while OTU 0010 proliferated under salt and fo conditions (**Fig. 5B**).

**Fig. 5:**
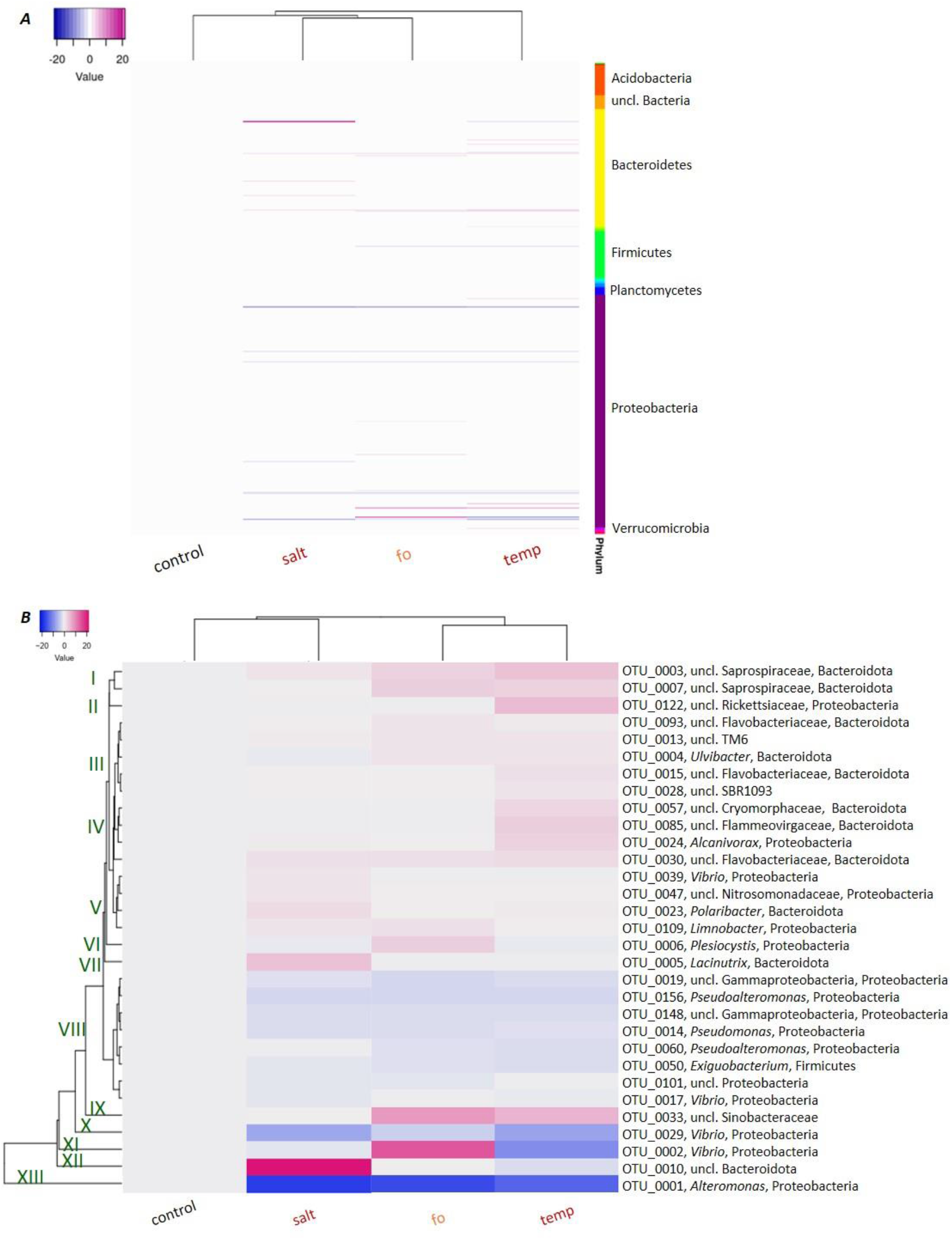
In-depth microbial community analyses. Heatmap and hierarchical clustering (based on the complete correlation of conditions) of OTUs in the microbiomes of polyps after 14 days of environmental challenge. Each column represents the mean of 6 replicates per treatment. Each row represents an OTU. The colors indicate fold changes of relative abundance compared to control conditions (blue, decreased abundance; red, increased abundance). **(*A*)** Holistic analysis of the identified 461 OTUs assigned to their phylum as indicated by the color code. **(*B*)** Heatmap and hierarchical clustering (cluster I – X) of most abundant OTUs (> 1 % relative abundance in control conditions) in the microbiomes.

Lastly, we combined all observations graphically in **Fig. 6**. Impaired fitness correlates with the increase of bacterial OTU clusters I, III, IV, V, VI, VII, IX, XI, XII and with the decline of OTU clusters VIII, X, XIII (**Figs. 5B, 6B**). The clusters VIII (8 OTUs), X (1 OTU), and XIII (1 OTU) comprise *Pseudolateromonas, Pseudomonas, Vibrio, Alteromonas*, and uncl. Gammaproteobacteria beside *Exiguobacterium*. Remarkably, the decrease of clusters X and XIII, each containing only one OTU (an *Alteromonas* and *Vibrio* species, respectively), correlates with an observed loss of host fitness (**Fig. 6**). Further, the increase of cluster I (2 OTUs of uncl. Saprospiraceae) and IX (1 OTU of uncl. Sinobacteraceae) were documented for all stress treatments (**Fig. 5B**). The remaining clusters produced a tendency to increase under stress conditions based on 2 of the three tested stress conditions (they decreased in the third). Notably, most OTUs classified as Bacteroidota were differentially affected by the treatments. Contrasting changes were also observed for four *Vibrio* OTUs: OTUs 0017 and 0029 became less abundant under all stress conditions, whereas OTUs 0002 and 0039 went up, depending on the condition. This observation points out that reporting findings at the genus level can be imprecise, and a more resolved analysis should be considered within complex microbiomes.

**Fig. 6:**
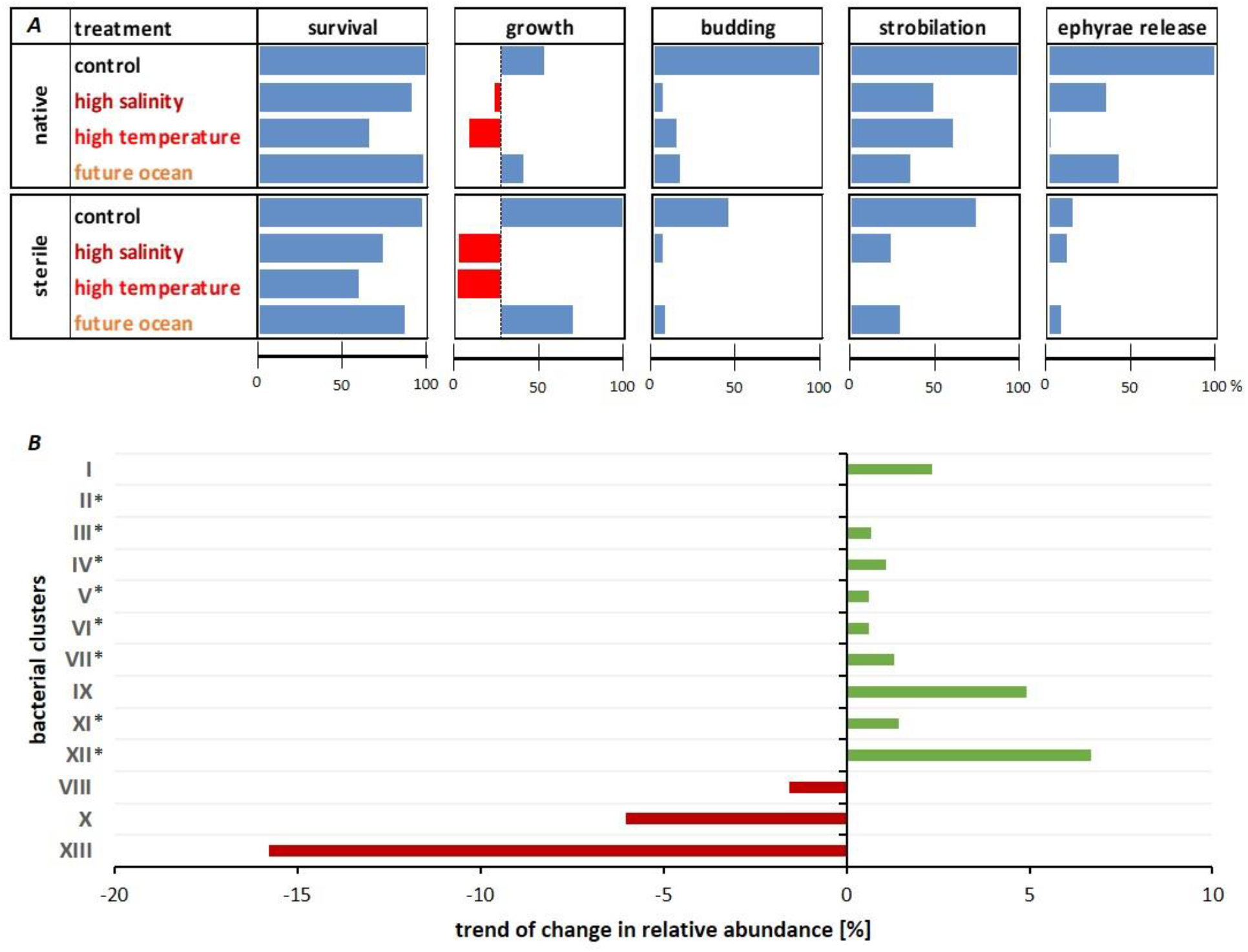
Impact of environmental changes on *Aurelia aurita* fitness traits in correlation to microbial presence/absence. (***A***) Fitness parameters. All data are expressed as % increase or decrease relative to control native animals taken as 100 %. Feeding rates are not shown as they did not significantly vary. (***B***) Trends of changes in relative abundances (%) in the clusters I to XIII of highly abundant OTUs as defined in Fig. 5B. Asterisks mark clusters with contrasting treatment-dependent changes, with bars showing the general trend for two out of three treatments.

## 4. Discussion

Over the past 100 years, the surface temperature has increased on average by 0.6 °C (Hansen, Sato & Ruedy 2022). Moreover, more frequent climatic extremes, like marine heatwaves, result in animal performance declines, mitigation, and local mortality (Oliver *et al*. 2018; Smith *et al*. 2022; Von Kietzell, Schurer & Hegerl 2022). In addition to temperature changes, historical records show that the ocean salinity increased by 4 % between 1950 and 2000 (Durack & Wijffels 2010). Many marine species are stenohaline, and their narrow range of salt tolerance limits their survival, reproduction, and germination (Evans & Kültz 2020). Salinity can act synergistically or antagonistically with other environmental stressors; for instance, salt stress was reported to cross-protect thermal stress (Przeslawski, Byrne & Mellin 2015; Velasco *et al*. 2019; Nahar *et al*. 2022). The capability of marine animals to adapt to future ocean scenarios is crucial for maintaining biodiversity and ecosystem functions (Baldassarre *et al*. 2022). Host-associated microbial communities represent a major factor regulating the host’s response to their external environment (Torda *et al*. 2017; Ziegler *et al*. 2017; Pita *et al*. 2018; Voolstra & Ziegler 2020; Baldassarre *et al*. 2022). Change in the composition of a host’s microbiome (both loss of taxa, shifts in relative abundance, or appearance of novel taxa) has been linked to adapted host fitness as a function of environmental change (Toby Kiers *et al*. 2010; Voolstra & Ziegler 2020). Correlative observational studies were reported for salt stress in algae (Ghaderiardakani, Quartino & Wichard 2020), thermal tolerance in sea anemones (Herrera *et al*. 2020; Randle *et al*. 2020), and heat stress in corals and sponges (Prazeres *et al*. 2017; Ziegler *et al*. 2017; Rubio-Portillo, Ramos-Esplá & Antón 2021; Santoro *et al*. 2021). Consequently, a shift in the microbiome toward a microbial community that supports host fitness could reinforce rapid host acclimatization (Marangon *et al*. 2021). High microbiome flexibility probably promotes metaorganismal acclimatization, at the risk of losing putatively essential associates and possibly allowing pathogen invasion (Voolstra & Ziegler 2020). Low microbiome flexibility in *Pocillopora* coral was linked to coral disease outbreaks, whereas high microbiome flexibility in *Acropora* corals was linked to rapid adaptation to escape the disease (Ziegler *et al*. 2019).

We consider the microbiome of *A. aurita* as crucial for host acclimatization, and we assume that the microbiota supports this host in adjusting to changes in the environment, such as temperature and salinity. A diverse and flexible microbiome might assist in maintaining host fitness in a climate-changed ocean. This assumption is supported by our host-fitness experiments conducted with sterile animals, which led to losses in survival, growth, and progeny output under standard (though sterile) conditions, exacerbated under environmental stressors. We had already demonstrated that bacteria function as a protective shield, and their absence impaired host fitness and affected life cycle decisions, resulting in the halt of offspring generation (Weiland-Bräuer *et al*. 2020a). Here, we demonstrate that the associated microbiota of *A. aurita* is changing in composition due to acute, sublethal temperature and salinity increases. This consequently affects host survival, and for those polyps that do survive, growth and asexual reproduction are impaired (**Fig. 2**). Note that energy-intensive fitness parameters related to reproduction were more affected than mere survival. Raising the salinity and temperature to sublethal levels impaired all analyzed fitness traits, leading to 66 % reduced survival rates and halting offspring generation. Several studies observed that environmental stressors diminish invertebrate reproduction (i.a., (Wang *et al*. 2018; Leger 2021)); however, only a few studies link microbiome shifts of these hosts to those effects (Carballo & Bell 2017).

We assume that changes in the microbial composition support acclimatization by the host, but drastic changes are associated with loss of microbial function, causing fitness deficits. In a natural setting, exposure to changed salinity or temperature may be more gradual than the abrupt changes applied here, possibly allowing for a slow but steady adaptation of the microbiome and its host. Nevertheless, during heatwaves, which are expected to increase in severity and frequency due to climate changes, local temperature and salinity changes can be relatively rapid, especially in shallow waters. When more moderate increases of salinity and temperature were combined in a future ocean scenario, this resulted in a less impaired fitness than for the more severe, single stressors (**Fig. 6**). This showed that the effects on host fitness correlate with the strength of the environmental stress, while salt-conveyed thermotolerance may also be involved. To our knowledge, salinity-conveyed thermotolerance in marine macroorganisms has only been described in corals (Ochsenkühn *et al*. 2017; Gegner *et al*. 2019) and data on *A. aurita* are lacking. Currently, it is unknown whether the bacterial community patterns and the response of the corals to different salinities are causally linked or whether they represent parallel responses of the host and its associated bacteria (Randle *et al*. 2020). Recent studies propose that osmolytes like floridoside might play a role in adjusting osmotic pressure by counteracting oxidative stress due to combined salinity and heat stress, thereby contributing to stress resilience (Ochsenkühn *et al*. 2017; Gegner *et al*. 2019). Similar studies would need to be conducted with *A. aurita* to gain deeper insights into the salinity-driven thermotolerance of this host.

Following analysis of the polyp’s microbiomes, we observed major changes in relative abundance that occurred on phylum, genus, and OTU levels (**Figs. 4, 5**). Highly-abundant genera like *Alteromonas, Pseudoalteromonas*, and *Pseudomonas* (all Proteobacteria) declined under all environmental stress conditions, while various unclassified genera assigned to Proteobacteria and Bacteroidota increased. Notably, some *Vibrio* OTUs increased, whereas others decreased (depending on the condition), indicating that reporting findings on genus level only can be imprecise. We demonstrate that approximately a quarter of the detected community members (26 %) maintain their relative abundance irrespective of environmental change. Other members may be interchangeable and act as microbiome regulators that maintain a constant microbiome functionality, irrespective of individual members during environmental change. Alternatively, those bacterial members that change in abundance due to environmental conditions may represent microbiome conformers that adapt to their surrounding environment and change the functionality of the complete microbiome (Ziegler *et al*. 2019). We noted that the 32 most abundant OTUs all changed their abundance as a result of environmental stress (**Fig. 5B**), and the intensity of the environmental stressor drives the degree of community change. Thermal tolerance of animals is assumed to be associated with an increase in Alpha- and Gamma-Proteobacteria (Webster *et al*. 2016; Pootakham *et al*. 2019; Baldassarre *et al*. 2022). That was not observed in our experiments, as the Proteobacteria phylum decreased under high salt or high temperature conditions. Alpha-Proteobacteria produce protecting antioxidants within the coral holobiont (Dungan *et al*. 2021), and Gamma-Proteobacteria representatives inhibited the growth of coral pathogens and provided additional nutrients for the host (Sabdono *et al*. 2015; Thompson *et al*. 2015). Clearly, such observations cannot be generalized and extended to different hosts, such as jellyfish. Impaired fitness of *A. aurita* polyps correlates with complex abundance shifts on the OTU level. We were able to link a decline of *Pseudolateromonas, Pseudomonas, Vibrio, Alteromonas, Exiguobacterium*, and uncl. Proteobacteria, and an increase of uncl. Saprospiraceae and uncl. Sinobacteraceae with impaired fitness. Primarily OTUs classified as Bacteroidota were affected by a specific stress treatment. Our results suggest that microbial communities are an essential factor affecting the response of animals to ambient temperature and salinity. The metaorganism concept should be considered for predicting species’ responses to global climate change.

## 5. Conclusions

The role of metaorganism’s microbiomes in host fitness and ecological interactions is increasingly evident. *A. aurita* is one of the main contributors to jellyfish blooms that cause enormous ecological and socioeconomic damage, and this study identifies the response of its microbiome to environmental challenges, coinciding with changes in the fitness of the polyps. A microbiome’s presence is essential for these animals’ stress tolerance, and microbial community changes correlate with impaired host fitness of *A. aurita* when the temperature or salinity is increased to sub-lethal levels. In a future ocean scenario, mimicked here by a combined but milder rise of temperature and salinity, the fitness of polyps was less severely impaired, together with condition-specific changes in the microbiome composition. Our results show that the effects on host fitness correlate with the strength of environmental stress, while salt-conveyed thermotolerance might be involved. Microbiome-mediated acclimatization and adaptation may provide a mechanism for hosts besides phenotypic plasticity. Thus, microbiome flexibility can be a fundamental strategy for marine animals to adapt to future ocean scenarios to maintain biodiversity and ecosystem functioning.

## Supplementary Materials

Tables S1-S6: PERMANOVA tests.: S1-Survival, S2-Growth, S3-Feeding rates, S4-Budding, S5-Segmentation (strobilation), S6-Ephyrae release

## Author Contributions

Conceptualization, N. W.-B.; methodology, N. P. and N. W.-B.; investigation, N. P. and N. W.-B.; formal analysis, N. W.-B.; bioinformatics analysis, S. G. and C.M. C.; data curation, C.M. C.; writing—original draft preparation, N. W.-B.; writing—review and editing, all authors; visualization, N. W.-B.; supervision, N. W.-B.; project administration, N. W.-B.; funding acquisition, N. W.-B. All authors have read and agreed to the published version of the manuscript.

## Funding

This work was conducted with the financial support of the DFG-funded Excellence Initiative “The Future Ocean” (CP1402).

## Acknowledgments

The authors thank Ruth A. Schmitz-Streit for providing laboratory space and appropriate equipment within the “Molecular Microbiology” working group of the Institute of General Microbiology, Kiel. We thank Sven Künzel and colleagues from the Department for Evolutionary Genetics of the Max Planck Institute for Evolutionary Biology for next-generation deep sequencing. Moreover, we thank the CRC1182 “Origin and Function of Metaorganisms” for network support.

## Conflicts of Interest

The authors declare no conflict of interest. The funders had no role in the study’s design, in the collection, analyses, or interpretation of data, in the writing of the manuscript, or in the decision to publish the results.

## Notes

### Competing Interest Statement

The authors have declared no competing interest.

