## Supplementary Materials for "Microbial community changes correlate with impaired host fitness of *Aurelia aurita* after environmental challenge"

**Title**

**PERMANOVA tests.**

Multivariate statistics were conducted using R for each fitness trait and all treatments (1-12). PERMANOVA p-values are depicted with their corresponding significance levels (*, < 0.05; **, < 0.01; ***,< 0.001; ns, not significant). Settings: Permutation test for adonis under reduced model; Permutation: free; No. of permutations: 9999 adonis2; native; sterile

**Table S1: Survival**

| Treatment | control | salt | temp | fo | control | salt | temp |
|---|---|---|---|---|---|---|---|
| salt | 0.001 *** | | | | | | |
| temp | 0.001 *** | | | | | | |
| fo | 0.265 ns | | | | | | |
| control | 0.266 ns | | | | | | |
| salt | | 0.001 *** | | | 0.001 *** | | |
| temp | | | 0.023 * | | 0.001 *** | | |
| fo | | | | 0.027 * | 0.001 *** | | |

**Table S2: Growth**

| Treatment | control | salt | temp | fo | control | salt | temp |
|---|---|---|---|---|---|---|---|
| salt | 0.001 *** | | | | | | |
| temp | 0.001 *** | | | | | | |
| fo | 0.001 *** | | | | | | |
| control | 0.001 *** | | | | | | |
| salt | | 0.001 *** | | | 0.001 *** | | |
| temp | | | 0.012 * | | 0.001 *** | | |
| fo | | | | 0.042 * | 0.002 *** | | |

**Table S3: Feeding rates**

| Treatment | control | salt | temp | fo | control | salt | temp |
|---|---|---|---|---|---|---|---|
| salt | 0.030 * | | | | | | |
| temp | 0.036 * | | | | | | |
| fo | 0.592 ns | | | | | | |
| control | 0.896 ns | | | | | | |
| salt | | 0.596 ns | | | 0.010 ** | | |
| temp | | | 0.498 ns | | 0.010 ** | | |
| fo | | | | 0.110 ns | 0.016 * | | |

### Table S4: Budding

| Treatment | control | salt | temp | fo | control | salt | temp |
|---|---|---|---|---|---|---|---|
| salt | 0.001 *** | | | | | | |
| temp | 0.001 *** | | | | | | |
| fo | 0.001 *** | | | | | | |
| control | 0.001 *** | | | | | | |
| salt | | 0.775 ns | | | 0.001 *** | | |
| temp | | | 0.001 *** | | 0.001 *** | | |
| fo | | | | 0.010 ** | 0.001 *** | | |

### Table S5: Segmentation (strobilation)

| Treatment | control | salt | temp | fo | control | salt | temp |
|---|---|---|---|---|---|---|---|
| salt | 0.001 *** | | | | | | |
| temp | 0.001 *** | | | | | | |
| fo | 0.001 *** | | | | | | |
| control | 0.001 *** | | | | | | |
| salt | | 0.002 ** | | | 0.001 *** | | |
| temp | | | 0.001 *** | | 0.001 *** | | |
| fo | | | | 0.007 ** | 0.001 *** | | |

### Table S6: Ephyrae release

| Treatment | control | salt | temp | fo | control | salt | temp |
|---|---|---|---|---|---|---|---|
| salt | 0.001 *** | | | | | | |
| temp | 0.001 *** | | | | | | |
| fo | 0.001 *** | | | | | | |
| control | 0.001 *** | | | | | | |
| salt | | 0.003 ** | | | 0.002 ** | | |
| temp | | | 0.897 ns | | 0.001 *** | | |
| fo | | | | 0.002 ** | 0.001 *** | | |
